# Efficient Synthesis of Mutants Using Genetic Crosses

**DOI:** 10.1101/359281

**Authors:** Aditya Pratapa, Amogh P. Jalihal, S. S. Ravi, T. M. Murali

## Abstract

The genetic cross is a fundamental, flexible, and widely-used experimental technique to create new mutant strains from existing ones. Surprisingly, the problem of how to efficiently compute a sequence of crosses that can make a desired target mutant from a set of source mutants has received scarce attention. In this paper, we make three contributions to this question.

First, we formulate several natural problems related to efficient synthesis of a target mutant from source mutants. Our formulations capture experimentally-useful notions of verifiability (e.g the need to confirm that a mutant contains mutations in the desired genes) and permissibility (e.g., the requirement that no intermediate mutants in the synthesis be inviable).

Second, we develop combinatorial techniques to solve these problems. We prove that checking the existence of a verifiable, permissible synthesis is **NP**-complete in general. We complement this result with three polynomial time or fixed-parameter tractable algorithms for optimal synthesis of a target mutant for special cases of the problem that arise in practice.

Third, we apply these algorithms to simulated data and to synthetic data. We use results from simulations of a mathematical model of the cell cycle to replicate realistic experimental scenarios where a biologist may be interested in creating several mutants in order to verify model predictions. Our results show that the consideration of permissible mutants can affect the existence of a synthesis or the number of crosses in an optimal one. Our algorithms gracefully handle the restrictions that permissible mutants impose. Results on synthetic data show that our algorithms scale well with increases in the size of the input and the fixed parameters.

## 1 Introduction

Engineering mutations in multiple genes is a powerful way to dissect cellular mechanisms, discover pathway structure, and understand the genetic basis of complex diseases. The genetic cross is a classical technique to create strains with mutations (e.g., knock-outs or overexpression) in multiple genes, especially in model organisms such as baker’s yeast, the fruitfly, mice, and *Arabidopsis thaliana*. Traditionally, biologists have created new strains by crossing existing mutants that they already have in the lab or can purchase easily. Strains carrying mutations in four, six, and even as many as 11 genes have been made in this manner [1, 20, 26]. More recently, developments in genome editing tools such as CRISPR-Cas9 have made it feasible to create mutations at multiple sites in a single experiment. This field is advancing very rapidly, with CRISPR-Cas9-like systems becoming available in several model organisms and with the development of new methods for multiplex gene editing [31].

Despite these advances, the genetic cross continues to remain a fundamentally useful experimental tool, including in combination with genome editing [24]. Since CRISPR-Cas9 edits require considerable efforts for verification, crosses are preferable if the mutants are already readily available. Moreover, in model organisms that have short generation times, performing a cross is often quicker than trying to edit two wild-type genes using CRISPR-based techniques. Therefore, in conjunction with high-throughput experimental techniques [6], genetic crosses open up the possibility of routine construction and characterization of mutants carrying alterations in several genes.

These observations inspire our research. We envision a scenario where a biologist has access in the lab to several source mutants, each carrying mutations in one or more genes; these mutants may have been created using a variety of techniques including crosses and genome editing. The biologist seeks to synthesize a target mutant using the genetic cross as the primary experimental tool.

### Our contributions

Our primary contributions are the definition of problems related to efficient synthesis of a target mutant from a set of source mutants using genetic crosses (Section 3) and combinatorial algorithms to solve these problems (Section 4). We first prove that when each source mutant contains mutations in at most three genes the problem of determining if there exists a synthesis of a target mutant is **NP** -complete. Next, we present a polynomial time algorithm to minimize the size of the synthesis (the number of crosses) of a target mutant when each source mutant contains mutations in at most two genes (Section 4.2). Furthermore, we provide a fixed-parameter tractable (FPT) algorithm for this problem with respect to the number of source mutants with three or more gene mutations (Section 4.4). Our final result deals with checking the existence of a permissible synthesis of a target mutant, i.e., ensuring that every intermediate mutant in the synthesis belongs to a permissible set of mutants. This problem is also **NP** -complete, in general. We provide a FPT algorithm with respect to the number of mutated genes in the target mutant (Section 4.5). This algorithm also minimizes the number of crosses.

These problems match an experimentalist’s growing repertoire of mutants. Initially, when there are only single gene mutants, the synthesis problem is trivial. Our first algorithm is useful when the experimentalist has made some double gene mutants. As triple mutants become available, the second algorithm is applicable, since it is FPT in the number of such mutants. When the set of available mutants increases further, the issue of permissibility becomes important, which our final algorithm addresses.

Two challenges confront us when we evaluate our algorithms: there is a dearth of prior literature on this problem (see Section 2) and there are no databases describing multi-gene (three or more) mutant strains that have been made in the lab. We use two complementary strategies to circumvent these gaps. First, we create a rich dataset of rescued and synthetic lethal target mutants by simulating a mathematical model of the cell cycle [15] for mutations in up-to-six genes (Section 5.1). We use mutants predicted to be viable by this model as permissible mutants. We apply our algorithms to compute syntheses for four-, five-, and six-gene target mutants. Permissibility has a material impact on a small fraction of mutants by either precluding the existence of a synthesis or by increasing the number of crosses needed in optimal syntheses. Second, we create synthetic datasets to assess the running times of our algorithms as functions of the fixed parameters (Section 5.2). Our results on these datasets show that the running time does grow with an increase in the value of the relevant fixed parameter.

## 2 Related Research

Mathematical models of cellular processes permit *in silico* simulations of multiple gene mutations [8,27,21, 28, 15]. Research on designing experiments based on model predictions has focused mainly on prioritizing the next-best or a small number of experiments, e.g., to distinguish between Boolean network models that fit the available data equally [12, 30, 5, 29, 3], to estimate the kinetic parameters of an ODE-based model [4, 17, 22], or to resolve model ambiguity [2, 11, 16, 18]. However, simulations of mathematical models or searching for informative experiments do not solve the problem of how to efficiently synthesize a strain carrying multiple mutations from existing strains.

This paper complements the only closely related research of which we are aware, which involves two of the current paper’s authors [25]. There, we defined the CrossPlan problem of optimizing the number of crosses needed in a verifiable, permissible synthesis of multiple target mutants. We improve upon the CrossPlan paper in three ways: (a) That paper used a model of a genetic cross that could produce all possible genotypes allowed by Mendelian inheritance. This type of genetic cross is experimentally challenging to implement since it requires verification of a number of genotypes that is exponential in the total number of mutations in the input mutants. Now, we propose a model of a genetic cross with a single output that harbors all the mutations in the two input mutant strains. This type of cross is experimentally more tractable and is usable in high-throughput workflows. (b) Earlier, we required that the input also contain a “genetic cross graph” that represents all possible ways of crossing source and permissible mutants. The worst case size of this graph can be quadratic in the number of source and permissible mutants. In contrast, in the current paper, we formulate problems that are more natural since they avoid the specification of the genetic cross graph in the input and the concomitant quadratic increase in the input size. (c) In the earlier paper, we proved that the synthesis problem was **NP** -complete and developed an integer-linear program (ILP) to solve it, but did not formally analyze the running time of the ILP. Here, we provide fixed-parameter tractable algorithms in natural parameters. In practice, we show that our algorithm for computing an optimal permissible synthesis is over 28 times faster than our previous approach (Section 5.2).

## 3 Definitions and Problem Formulations

A *mutant μ* is equivalent to a set of genes, each of which has been mutated in some manner experimentally, e.g., a deletion (knock-out), disruption, point mutation, over-expression, etc. We will use the term *representative set* to denote the set of genes deleted in a mutant. An element of such a set represents the mutated gene as well as any other necessary experimental information, e.g., the name of the selectable marker introduced with a gene deletion (in budding yeast; see below) or the recessive deleterious mutations and balancers that accompany a mutation (in the fruitfly).

A *cross* is an operation that takes two mutants as input and produces another mutant as output. If *μ*_1_ and *μ*_2_ are the inputs to a cross with the corresponding representative sets *R_1_* and *R_2_*, then the output is a mutant whose representative set is *R_1_* ∪ *R_2_*. We say that a cross is *verifiable* if the representative sets of the two input mutants to the cross are disjoint.

We formulated these definitions based on experimental and biological considerations. In many organisms, e.g., budding yeast, each gene deletion construct in a mutant strain must have a unique selectable marker that can confirm the deletion, e.g., *a∆::kanR* represents a deletion of gene *A* and its replacement by the *kanR* G418 resistance gene. Only strains with gene *A* deleted will grow in the presence of the an tibiotic G418, which normally kills budding yeast cells. Similarly, *bΔ::nat* represents a strain carrying a deletion of gene *B* that will grow in the presence of the antibiotic nourseothricin. An experimenter crossing these mutants can select for the *a∆b∆* double mutant by growing cells in the presence of both G418 and nourseothricin. Thus, markers or other strategies involving recessive mutations and balancers enable the experimenter to ignore any strain produced by the cross whose representative set is a subset of *R_1_* U *R_2_*. Moreover, the need to verify only one output strain for each genetic cross facilitates high-throughput experimental workflows. We based our definition of a genetic cross on this rationale. Note that in the example above, the selection of the *aΔbΔ* double mutantis successful only if the two strains carry different markers. Hence, the notion of verifiability of a cross captures the natural requirement that when we cross two mutants, each gene in a representative set is (implicitly) associated with a marker that is not associated with any other gene in that or the other mutant. By abstracting away unimportant experimental details, these definitions cleanly capture the essential features of a cross while also leading to challenging computational problems.

Given a set of source mutants *S* and a set *T* of target mutants, we say that a sequence of crosses ⟨ *x*_1_, *x _2_*,…, *x_k_*⟩ is a *synthesis of T from S* if (i) for every mutant *μ* ϵ *T*, there is a unique cross *x_i_* such that *μ* is the output of *x_i_*, (ii) both the inputs to *x_1_* are from *S* and (iii) for every *i*, 2 ≤ *i ∆ ≤ k*, each input to *x_i_ i*is either a member of *S* or an output of one of the crosses *x*_1_, *x*_2_,…, *x*_i_–1.

The *size* of a synthesis is the number of crosses it contains. We say that a synthesis is *verifiable* if every cross in it is verifiable. In such a synthesis, we say that a mutant is *intermediate* if it is not in *S* U *T* and is used as the input to another cross in the synthesis. Given a set *P* of mutants, we say that a synthesis is *permissible with respect to P* if every intermediate mutant in the synthesis is in *P*. The notion of permissibility takes inspiration from applications where we are interested in computing syntheses that avoid particular intermediate mutants, e.g., when we know that mutants involving a certain gene are certain to be lethal or when mathematical models predict that some mutants are likely to be inviable. Without loss of generality, we assume that a synthesis is *minimal,* i.e., for every cross *x_i_*, 1 ≤*i ≤ k,* where the output of *x_i_* is not a member of *T*, this output is used as an input to a subsequent cross *x_j_*, *i* ≤ *j* ≤ *k*. Figure 1 illustrates several of these definitions.

**Figure 1:**
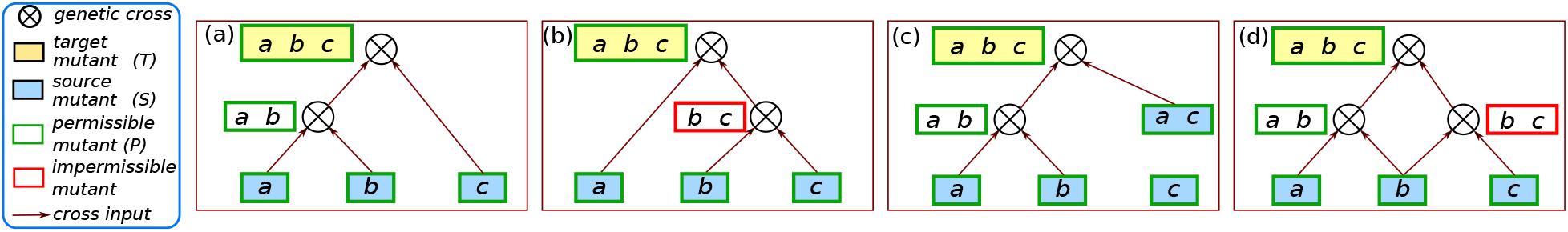
Illustration of different types of synthesis. Each rectangle denotes a mutant. Each figure shows a synthesis of *{a, b, c}* from source mutants (blue). The rectangle next to a cross denotes the output mutant of that cross. (a) A verifiable, permissible synthesis. (b) A verifiable synthesis that is not permissible because the intermediate mutant *{b, c}* is not permissible. (c) A permissible synthesis that is not verifiable since the two input mutants to the second cross both have gene a in their representative sets. (d) A synthesis that is neither verifiable nor permissible.

We now formulate the main problem that we study followed by several important special cases:

### Minimum Verifiable and Permissible Synthesis (VPS)

Given a set *S* of source mutants, a target mutant *μ*, and a set *P* of intermediate mutants, compute a verifiable synthesis of *μ* of smallest size from *S* that is permissible with respect to *P*, provided such a synthesis exists.

**VS** The inputs to this problem are the source set *S* and the target mutant *μ*. We seek to compute a smallest synthesis of *μ* starting from *S*. In other words, this problem is identical to VPS except that all mutants are permissible.

**VS-2** This version of VS assumes that the representative set of each mutant in *S* has at most two elements. This problem simulates a scenario where an experimentalist has several single-gene mutants and has used them to make double-gene mutants as well.

**VS-FL** This version of VS assumes that a fixed number *q* of mutants in *S* whose representative sets are of size three or more (i.e., *q* is a constant independent of the problem size). The name is an abbreviation for “VS with a Few Large representative sets.” Here, the experimentalist also has access to a small number of mutants with mutations in three or more genes. We are also interested in determining whether a synthesis with the appropriate properties exists. We will use **EVS** to denote the existence version of VS.

## 4 Algorithms for Verifiable and Permissible Synthesis

We first consider the VS problem and its variants. In Section 4.1, we show that EVS is **NP** -complete even when each source mutant has at most three deleted genes. It follows that VS is **NP** -hard, in general. Next, we show that VS-2 can be solved in polynomial time (Section 4.2). Using this algorithm as a sub-routine, we establish that VS-FL is FPT, where the parameter is the number of source mutants with three or more deleted genes (Section 4.4). Finally, we develop a FPT dynamic programming algorithm for VPS, where the parameter is the largest size of a representative set of the target mutant (Section 4.5).

We assume that all the problems take as input a starting set *S* of *s* mutants, specified by representative sets *R_1_*, *R_2_*, …, *R _s_* and a target mutant *μ’* with the representative set *R’*, where *r* = | *R’*|. We assume without loss of generality that 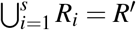 since any mutant with a representative set containing a gene that is not in *R’* may be removed from *S*. For the VPS problem, the set of permissible mutants is denoted by *P*.

### 4.1 EVS is NP-complete

We first recall the definition of EVS.

#### Existence of a Verifiable Synthesis (EVS)

##### Instance

A set *S* = { *μ*_1_,*μ*_2_,…,*μ_s_*} of s source mutants, the representative set *R_i_* for each mutant *μ_i_*, 1 ≤ *i* ≤ *s*; the representative set *R’* for a target mutant *μ’*.

##### Question

Is there a verifiable synthesis of *μ’* from the given set *S* of mutants?

To prove the **NP** -completeness of EVS, we use a reduction from the following problem.

#### Exact Cover by 3-Sets (X3C)

##### Instance

A ground set *Y =* {*y*_1_,*y*_2_,…,*y_3n_*} of 3*n* elements for some positive integern, a collection *Z* = {*Z*_1_,*Z*_2_,…,*Z_m_*}, where *Z_j_* ⊆ *Y* and |Z_j_| | = 3,1 ≤ *j* ≤ *m*.

##### Question

Is there a subcollection *Z’* ⊆ *Z* such such that | *Z’*| = *n* and 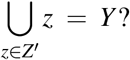

It is well known that X3C is **NP** -complete [10]. Note that in the specification of X3C, the ground set *Y* has 3*n* elements, the required subcollection *Z^’^* is of *size n* and each subset in *Z^’^* has exactly three elements. Therefore, when there is a solution *Z^’^* to an X3C instance, the subsets in *Z^’^* are pairwise disjoint.

###### Theorem 4.1. EVS is NP-complete even when the representative set for each source mutant has at most three elements

*Proof*. It is easy to see that EVS is in **NP** since one can guess a subset *S’* of *S* and verify in polynomial time that the sets in *S’* are pairwise disjoint and that their union is equal to *S’*, the representative set of the target mutant *μ’*.

To prove **NP** -hardness, we use a reduction from X3C. Let an instance of X3C be specified by the set *Y* (with 3n elements) and the collection *Z* of *m* 3-element subsets of *Y*. We construct an instance of EVS as follows.

1. The set *S =* { *μ*_1_, *μ* _2_,…, *μ_m_*} of available mutants is in one-to-one correspondence with the collection of sets *Z* of the X3C instance. For mutant *μj,* the representative set is *Zj*, 1 ≤ *j* ≤ *m*. Thus, the number of source mutants is *m* and the representative set for each input mutant has exactly three elements.

2. The representative set *S^’^* for the target mutant *μ’* is *Y*.

It is easy to see that this construction can be carried out in polynomial time.

Suppose there is a solution 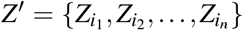 to the X3C instance. A verifiable synthesis of *μ’* consisting of the sequence ⟨ *x*_1_, *x*_2_,…, *x_n—1_*⟩ of crosses is as follows:

a. The inputs to *x*_1_ are mutants 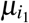 and 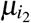
b. For each *j*, 2 ≤ *j* ≤ *n* — 1, the inputs to *x_j_* are the mutant 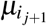 and the output of *x_j_*−1.

Since the union of the sets in *Z’* is equal to *Y*, the output of *x_n-1_* is the mutant *μ’*. Further, since the sets in *Z’* are pairwise disjoint, this is a verifiable synthesis. In other words, we have a solution to the EVS instance.

Suppose we have a solution to the EVS instance. Let 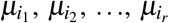 denote the source mutants used in the synthesis and let 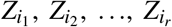 denote their respective representative sets. Thus, 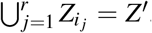. Moreover, since we have a verifiable synthesis of *μ’*, each pair of representative sets used in the synthesis is disjoint. Further, each representative set has exactly three elements and their union is the set *Z’*, where |*Z’*| = 3*n*. Thus, the number of representative sets used must be exactly *n*. In other words, the sets 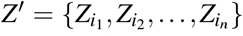 constitute a solution to the X3C instance, and this argument completes the proof of Theorem 4.1.⃞

### 4.2 An Efficient Algorithm for VS-2

Here, we consider VS-2, a restricted version of EVS, where the representative set of each input mutant has *at most two* elements. We show that VS-2 can be solved efficiently, thus providing a clear delineation between the **NP** -hard and efficiently solvable versions of EVS. Our polynomial time algorithm for VS-2 relies on a known algorithm for the following problem.

#### Degree Constrained Subgraph (DCS)

Given an undirected graph *G* (*V, E*) with | *V*| = *n*, and non-negative integers a _i_ and *b*_i_ for each node v_i_ ϵ *V*, 1 ≤ *i* ≤ *n*, is there a subgraph *G’*(*V, E’*) of G, where *E’* ⊆ *E*, such that for each node *v_i_*, 1 ≤ *i* ≤ *n*, the degree of v_i_ in *G’*, denoted by *d’*(*v_i_*), satisfies the condition *a _i_* ≤ *d’(v _i_*) ≤ *b_i_*?

The DCS problem can be solved in time 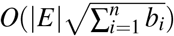 [9]. Moreover, this algorithm can be modified to compute a subgraph with the fewest edges that satisfies the conditions of the problem with the same running time [13]. We obtain our result for VS-2 by reducing it to DCS and using the algorithm for DCS in [13]. The details of our algorithm for VS-2 appear in Algorithm 1. See Figure 2 for an illustration.

**Figure 2:**
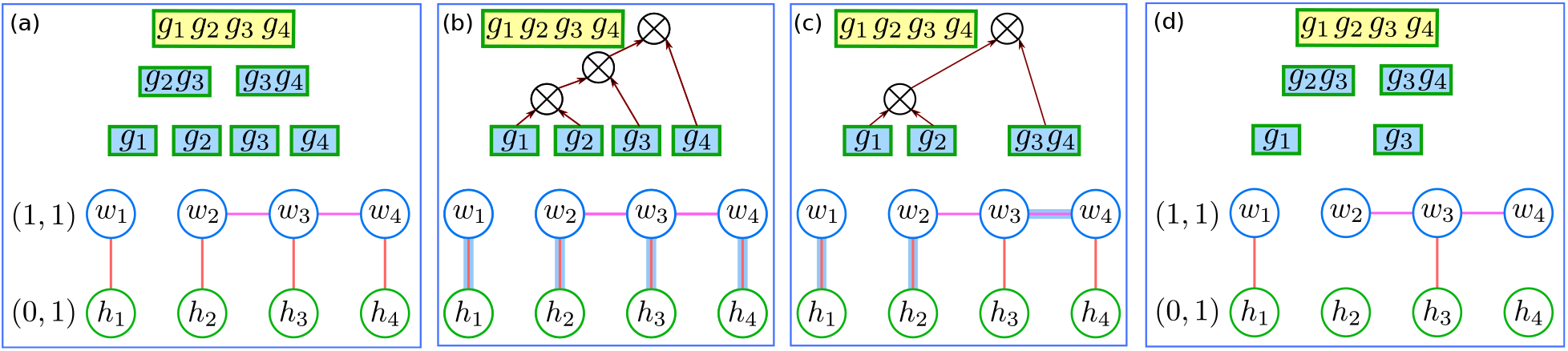
Reduction of VS to DCS. (a) The top show the input to VS with six source mutants (blue boxes) and one target mutant (yellow box). The bottom shows the corresponding DCS instance. Numbers in parentheses indicate bounds on degrees. (b) Feasible solution to DCS (edges with a blue outline). This solution is equivalent to three crosses to make the target mutant (*g*_1_, *g*_2_, *g*_4_, *g*_4_). (c) An optimal solution to DCS that yields two crosses to make (*g*_1_, *g*_2_, *g*_4_, *g*_4_). (d) An instance of VS that does not have a solution. Such an input may arise when single gene deletions of *g*_2_ and *g*_4_ make the cell inviable but the double deletions (*g*_2_, *g*_3_) and (*g*_3_, *g*_4_) are obtained via transformation.

We now establish the correctness of the algorithm and estimate its running time to prove the following theorem.

##### Theorem 4.2

*Algorithm 1 correctly solves the* VS-2 *problem in* 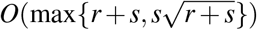 *time using O*(*r* + *s*) space, where s is number of source mutants and r is the size of the representative set of the target mutant.

*Proof*. We will refer to the steps of this algorithm throughout this proof.

**Correctness**: We first consider the case where the algorithm produces a solution and prove that the representative sets of the mutants in the solution are pairwise disjoint and their union is the set *R’*. Let 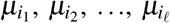, be the mutants chosen in the solution; their respective representative sets are 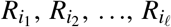. Suppose there is a pair of representative sets in the solution, say 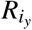 and 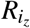, which are not disjoint. Let *g_p_* be an element that appears in both 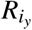 and 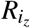. Since each representative set represents an edge in the DCS instance, it follows that the degree of the node *w_p_* ϵ *V*_1_ corresponding to *g_p_* is at least two. This contradicts the requirement that the upper bound on the degree of each node in V_1_ is one. Thus, the representative sets in the solution are indeed pairwise disjoint. Now suppose that 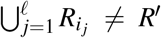. Thus, some element g _q_ ϵ *R’* does not appear in any of the sets in the solution. In other words, in the solution to the DCS instance, the degree of node *w_q_* ϵ *V*_1_ corresponding to *x_q_* is zero. This contradicts the requirement that the lower bound on the degree of each node in *V*_1_ is one. Thus, whenever the algorithm produces a solution, the chosen mutants can be used to construct a verifiable synthesis of the required mutant *μ’*.

**Algorithm 1:**
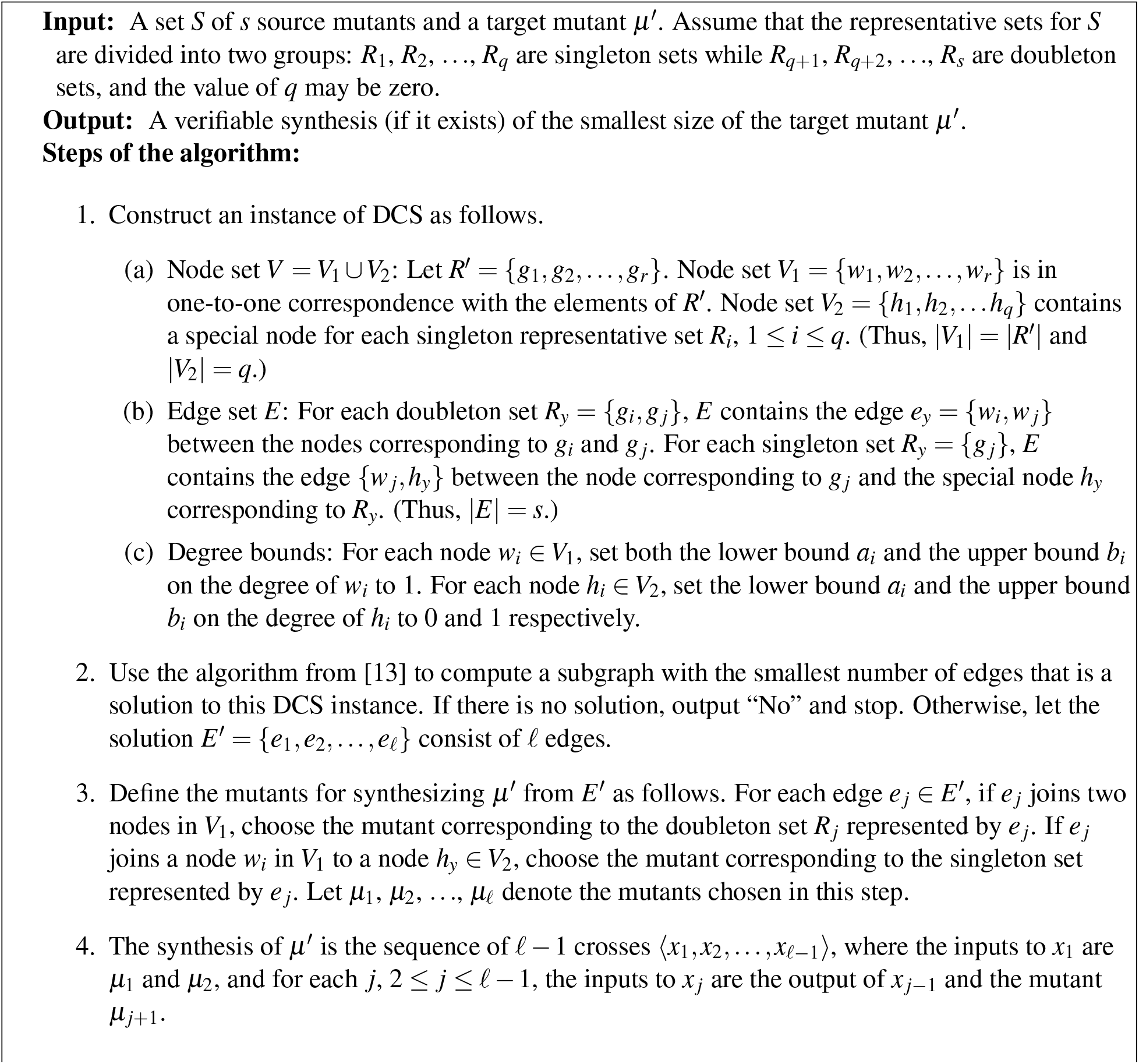
An algorithm for VS-2.

Now, suppose the algorithm outputs “No”. We show by contradiction that there is no solution to the VS-2 instance. Assume to the contrary that there is indeed a solution with mutants 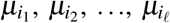 with corresponding representative sets 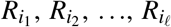. Since each of these sets is represented by an edge in the DCS instance, let 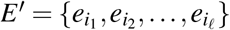 represent the corresponding edges in G. We argue that the subgraph *G’(V, E*’) is a solution to the DCS instance by considering each node in *V*. There are two cases depending on whether the node is in *V*_1_ or in *V*_2_.

<u>Case 1:</u> Consider any node *w_p_* ϵ *V _1_* We need to show that the degree of *w_p_* in *G’* is 1. To see this, note that node *w_p_* appears in some edge of *E*’ since 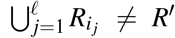. Thus, the degree of *w_p_* in *G’* is at least 1. If degree of *w_p_* is ≥ 2, then the corresponding element *x_p_* appears in two or more sets of the solution; this contradicts the assumption that the solution sets are pairwise disjoint. Thus, the degree of *w_p_* is 1.

<u>Case 2:</u> Consider any node *h_q_* ϵ *V*_2_. The degree of *h_q_* in *G’* is at most 1 since each node *h_q_* in *V*_2_ has a degree of 1 in *G*.

Thus, we contradict the assumption that the algorithm outputs “No”. This argument completes the proof of correctness.

**Analysis of running time and space used.** The constructed DCS instance has *r* + *q* nodes (where *r =* | *R’*| and *q* ≤ *s* is the number of singleton representative sets of source mutants) and *s* edges. Thus, we can construct the graph in *O* (*r* +*s*) time. The upper bound on the degree of each node is 1. Thus, we can obtain the optimal solution to the DCS instance in 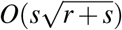 time [9, 13]. Since the solution produced by the algorithm for DCS has at most s edges, we can find the representative sets in the solution in *O* (*s*) time. Thus, the overall running time of the algorithm is 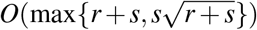. Further, since the size of each representative set of a source mutant is at most two and that of the target mutant is *r,* the space used by the algorithm is *O* (*r* + s). This discussion completes the proof of the theorem.⃞

### 4.3 An ILP for the DCS Problem

In Section 4.2 we showed that VS-2 can be solved in polynomial time by a reduction to the DCS problem. In this section, we develop a simple {0,1} integer linear program (ILP) to solve the DCS problem. We use this ILP in lieu of Algorithm 1 in our experimental results in Section 5 since our problem instances are reasonably small in practice.

Consider a DCS problem instance specified by a graph *G(V,E*), where *V =* {*v*_1_, *v*_2_,…, *v_n_*} and *E =* {*e_1_*, *e*_2_,…, *e_r_*}. Let the lower and upper bounds on the degree of node *v_i_* be *a_i_* and *b_i_* respectively; that is, in the subgraph *G’(V,E’)* to be found, degree of *v_i_* must satisfy the condition *a_i_*≤ degree of *v_i_* ≤ *b_i_*,1 ≤ *i* ≤ *n*. We also want to minimize the number of edges in *E^’^*.

The ILP has a {0,1} variable *x_i_* for each edge *e_i_*, 1 ≤ *i* ≤ *r,* with the following interpretation: if *x_i_* = 1, edge *e_i_* is included in the solution; otherwise, *e_i_* is not in the solution. So, the objective of the ILP is:

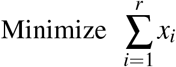

For each node *v_i_,* there are two constraints, corresponding to the upper and lower bounds on the degree of *v_i_* in *G’*. Let *Q_i_* ⊆ *E* be set of edges of *G* which are incident on node *v_i_*. The constraints for node *v_i_* are:

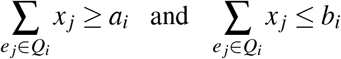

Thus, we get a total of 2*n* constraints. For any node *v_i_* whose the lower bound *a_i_* is 0, we omit the corresponding constraint. We solve this ILP using IBM CPLEX software.

**Algorithm 2:**
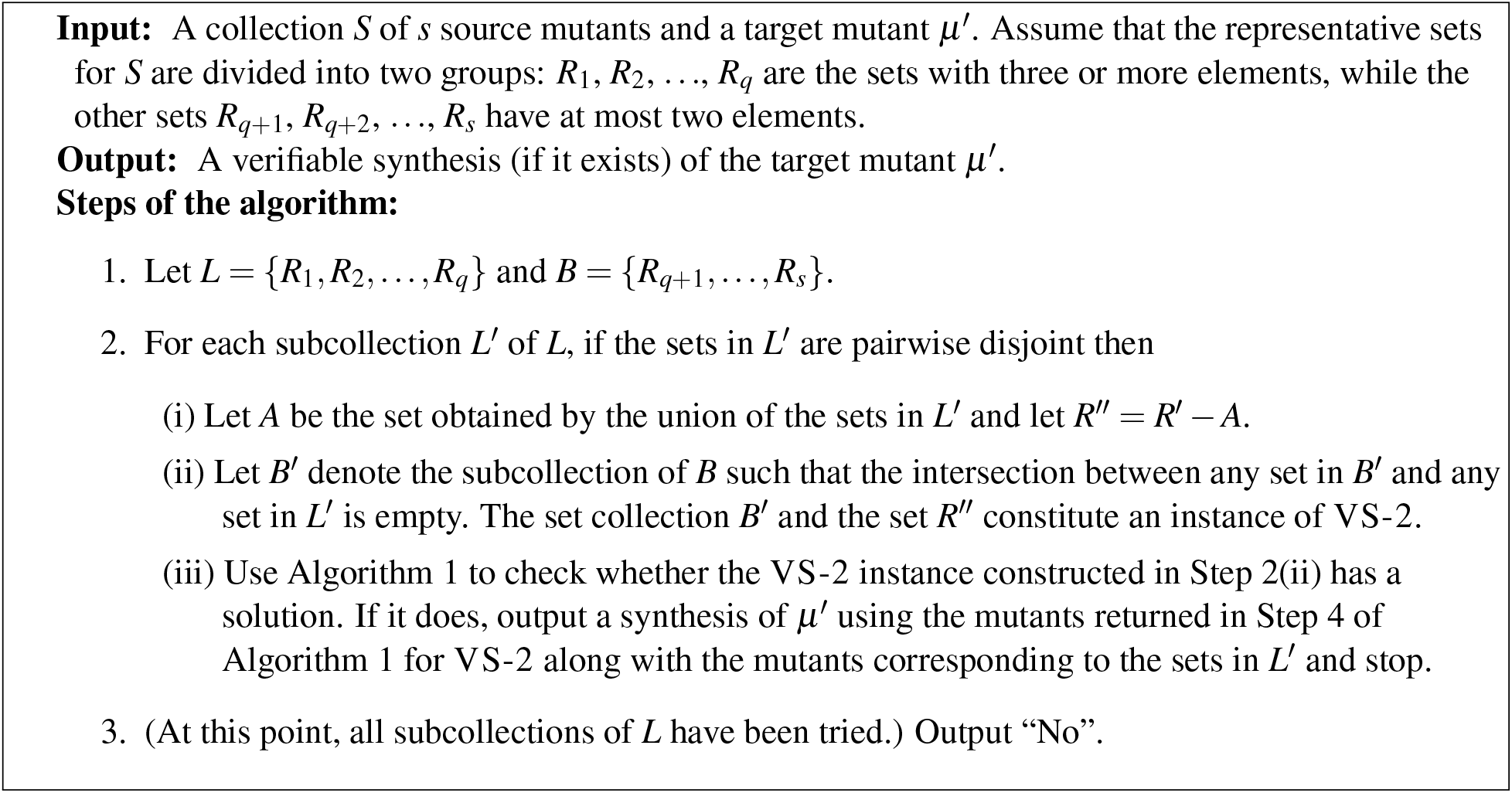
An algorithm for VS-FL.

### 4.4 Fixed Parameter Tractability of VS

Here, we consider VS-FL, another restricted version of VS, where the number *q* of mutants whose representative sets are of size 3 or more is *fixed* (i.e a constant independent of the problem size). We assume that *q >* 0; otherwise, we have the VS-2 problem which can be solved in polynomial time (Section 4.2). We show that VS-FL can be solved in 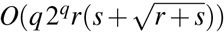 time using *O* (*rs*) space, where *r* = |*R’*| and s is the number of source mutants. More importantly, this result points out that VS-FL is *fixed parameter tractable,* with respect to the parameter *q* [19], since the component of the running time that is independent of *q* is polynomial in the input size.

The principle underlying our algorithm for VS-FL is simple. Let *L* denote the collection of representative sets (of source mutants) whose size is at least three. We use *q* to denote the fixed size of *L*. We consider all the possible 2^q^ subcollections (including the empty subcollection) of *L*. For each such subcollection *L’*, the union of the sets in *L^’^* contributes some of the elements of the representative set *R’* of the target mutant *μ’*. The remaining elements of *R’* must be synthesized using mutants whose representative sets are of size at most two, giving rise to an instance of the VS-2 problem, which can be solved using the algorithm of Section 4.2. The details of our method appear in Algorithm 2. Note that this algorithm determines if a verifiable synthesis of *μ’* exists. It is easy to modify it to compute a verifiable synthesis of smallest size: In Step 2(iii), instead of stopping when we find a synthesis, we record the size of the synthesis and return a smallest synthesis computed over all executions of this step.

We now prove the following theorem.

#### Theorem 4.3.

*Algorithm 2 solves the* VS-FL *problem in* 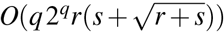*time using O(rs) space. Thus, the* VS-FL *problem is fixed parameter tractable in q, the number of representative sets of input mutants with three or more elements*.

*Proof*. We prove the theorem by establishing the correctness of Algorithm 2 and then showing that its running time has the appropriate form to imply the fixed parameter tractability of VS-FL.

**Correctness:** It is obvious that when the algorithm returns a solution, the representative sets in the solution are pairwise disjoint and their union is equal to *R’*, the representative set of the target mutant *μ’*. So, we need only consider the case when the algorithm outputs “No”. For the sake of contradiction, assume that the algorithm outputs “No”, yet there is a solution to the VS-FL instance. Suppose the solution uses a subcollection *L’* of *L* (where *L’* may be empty) and subcollection *B’* of *B* = *{R_q+1_*,…,*R_s_*}. Since the algorithm considers all subcollections of *L* (including the empty subcollection), it considers *L ^’^* at some stage. The sets in the solution are pairwise disjoint and the union of the sets in *B’* ∪ *L’* is equal to *R’*. Let the union of the sets in *L’* be denoted by A. Thus, the union of the sets in *B’* must be equal to *R’’* = *R’ — A*. Note that each set in *B’* has size at most two. Hence, the subcollection *B’* and set *R’’* constitute an instance of VS-2 for which a solution is B’. Therefore, the algorithm for VS-2 cannot output “No”. This contradiction completes the correctness proof.

**Analysis of running time and space used:** Since | *L*| = *q*, the number of iterations of the for loop in Step 2 of Algorithm 2 is at most 2*^q^*. Let | *R’*| = *r*. Thus, each representative set has size at most r. For each subcollection *L’* (which has at most *q* sets), the set *R’’* in Step 2(i) of Algorithm 2 can be computed in *O* (*rq*) time. Further, the subcollection *B’* of *B* = {*R_q_*+1,…,*R_s_*} such that the intersection between any set in *B’* and any set in *L’* is empty can be constructed in *O(qrs)* time. The resulting instance of VS-2 has at most *s* sets (each of size one or two) and a representative set *R’’* of size at most *r*. Therefore, by Theorem 4.2, the VS-2 instance can be solved in time 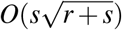 time. Thus, the running time for each subcollection *L’* is 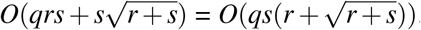. Since the algorithm tries at most 2*^q^* subcollections, the overall running time is 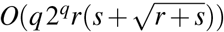. This running time has the form *O(h(q)N^O^*^(1)^), where *h(q)* = *q2^q^*depends only on the parameter *q* and *N = rs* is the size of the VS-FL instance. Hence, VS-FL is fixed parameter tractable with respect to the parameter *q*. Further, since the size of the representative set of each mutant is at most *r* and the total number of mutants (i.e., source and target mutants) is s + 1, the space used by the algorithm is *O*(*rs*). This completes the proof of the theorem.⃞

### 4.5 Fixed Parameter Tractability of VPS

We now turn our attention to the VPS problem. Recall that we are given a set *S* of *s* source mutants, a set *P* of *p* permissible mutants, and a target mutant *μ’*. Our goal is to compute a verifiable synthesis of *μ’* of smallest size from the mutants in *S* that is permissible with respect to *P,* if such a synthesis exists. We present an FPT algorithm for this problem in the parameter *k* = | *R’*|, where *R’* is the representative set of the target mutant *μ’*.

We assume that every mutants in *S* and in *P* has a representative set of size at most *k,* since only such a mutant can be used in the synthesis of *μ’*. We also assume that each source mutant is permissible, as is the target mutant. We start with a simple definition that captures the notion of verifiable and permissible synthesis.

**Algorithm 3:**
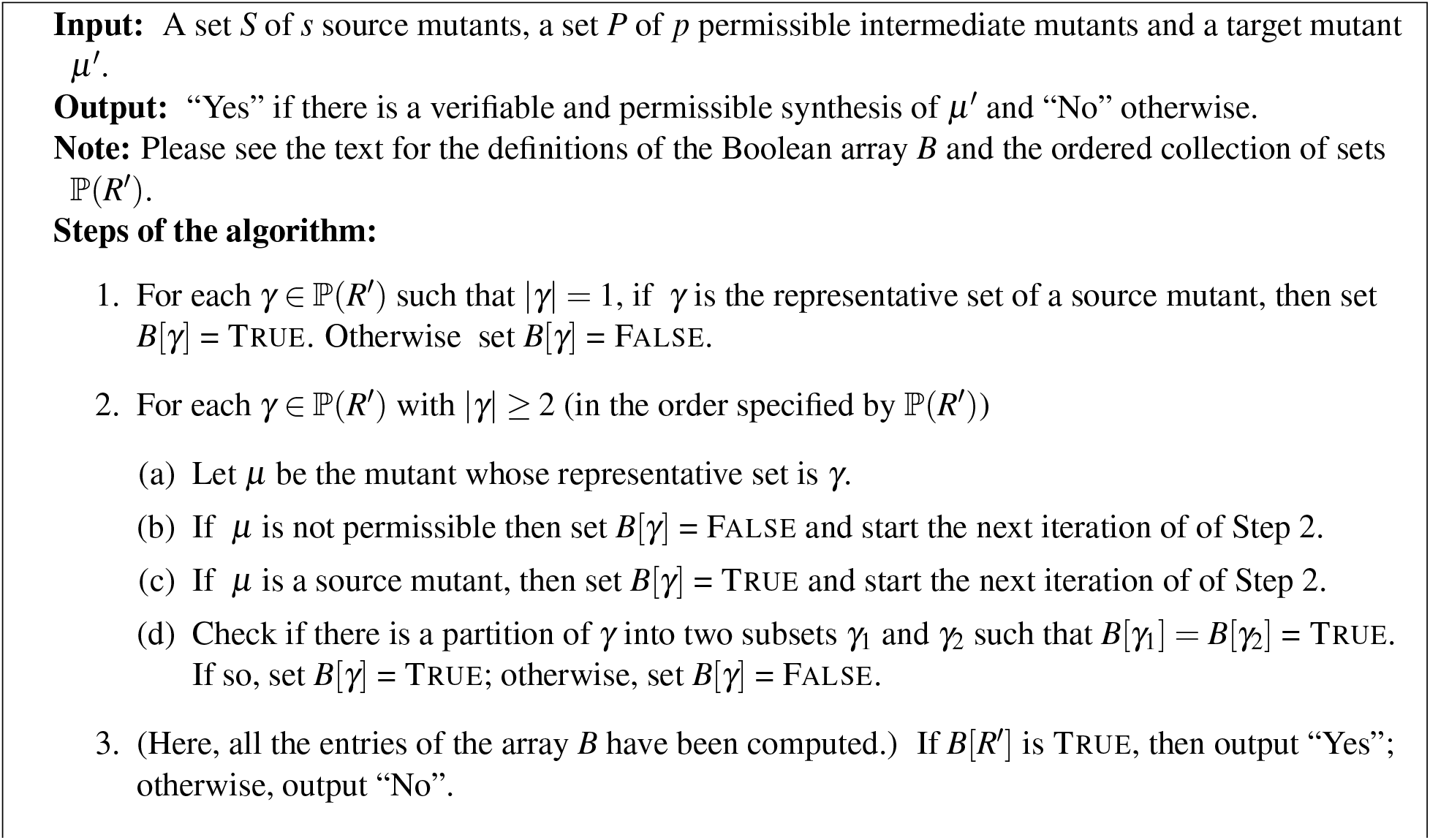
A dynamic programming algorithm for EVPS.

#### Definition 4.1

*A target mutant μ’ is* P-realizable *(for “Physically realizable “) if (α) μ’ is a source mutant or (b) there is a verifiable synthesis of μ’ from the source mutants such that each intermediate mutant in the synthesis is permissible*.

Thus, our goal is to determine whether a given target mutant *μ’* is P-realizable. In developing our algorithm, it is useful to define a computational notion of realizability, which we call C-realizability. A recursive definition of this notion is as follows.

#### Definition 4.2

*A target mutant μ’ with representative set R’ is* C-realizable *(for “Computationally realizable”) if (a) μ’ is a source mutant or (b) there is a partition of R’ into two subsets* 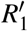 *and* 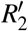 *such that the two corresponding mutants* 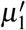 *and* 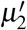 *are both permissible and C-realizable*.

The following theorem states that the notions of P-realizability and C-realizability are equivalent.

##### Theorem 4.4

*A permissible target mutant μ’ is C-realizable if and only if it is P-realizable*.

*Proof*. **Part 1 (If):** Suppose *μ’* is permissible and P-realizable. We use induction on the size of *R’*, the representative set of *μ’* to show that *μ’* is C-realizable. Let *α ≥* 1 be the minimum size of the representative set of a source mutant. Since *μ’* is permissible, | *R’*| ≥*α*.

**Basis:** |*R’*| = *α*. In this case, since *μ’* is permissible, it is a source mutant. By definition, each source mutant is C-realizable. Hence, the basis holds.

**Inductive hypothesis:** Assume that for some *r* ≥ *α*, every permissible and P-realizable mutant *μ* whose representative set is of size ≤ *r* is also C-realizable.

**Inductive proof:** Let *μ’* be a permissible and P-realizable mutant whose representative set *R’* is of size *r* + 1. We need to prove that *μ’* is also C-realizable.

If *μ’* is a source mutant, then it is C-realizable by definition. So, assume that *μ’* is not a source mutant. Thus, there is a verifiable synthesis of *μ’* such that each intermediate mutant in the synthesis is permissible.

Consider the last cross operation x in this synthesis. Let 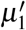 and 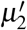 denote the mutants which form the inputs to *x*. Hence, both 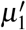 and 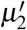 are permissible and the corresponding representative sets 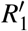 and 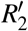 are disjoint. Thus, there is a verifiable synthesis of 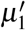 and 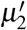 such that each intermediate mutant in the synthesis is permissible. In other words, both 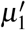 and 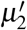 are permissible and P-realizable. Further, the sizes of the corresponding representative sets 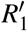 and 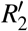 are ≤ *r*. Therefore, by the inductive hypothesis, both 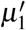 and 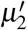 are C-realizable. In addition, sets 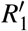 and 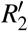 are disjoint and their union is *R’*. In other words, 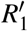 and 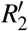 form a partition of *R’* and the two corresponding mutants 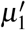 and 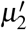 are both C-realizable. Thus, by definition, *μ’* is also C-realizable. This completes the proof of Part 1.

**Part 2 (Only If):** Suppose *μ’* is permissible and C-realizable. We again use induction on the size of *R’* to show that *μ’* is also P-realizable. As before, let *α* ≥ 1 be the minimum size of the representative set of a source mutant. Since *μ’* is permissible, | *R’*| ≥ *α*.

**Basis:** | *R’*| = *α*. In this case, since *μ’* is permissible, it is a source mutant. By definition, each source mutant is P-realizable. Hence, the basis holds.

**Inductive hypothesis:** Assume that for some *r* ≥ *α*, every permissible and C-realizable mutant *μ* whose representative set is of size ≤ *r* is also P-realizable.

**Inductive proof:** Let *μ’* be a permissible and C-realizable mutant whose representative set *R’* is of size *r* + 1. We will prove that *μ’* is also P-realizable.

If *μ’* is a source mutant, then it is P-realizable by definition. So, assume that *μ’* is not a source mutant. Since *μ’* is C-realizable, there is a partition of *R’* into two two subsets 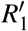 and 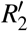 such that the two corresponding mutants 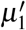 and 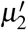 are both permissible and C-realizable. Note that 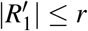 and 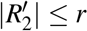. Thus, by the induction hypothesis, both 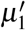 and 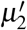 are permissible and P-realizable. Now, construct a synthesis of *μ’* by adding a new cross operation x whose inputs are the mutants 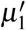 and 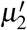. (Note that these may be source mutants or outputs of cross operations.) It is easy to see that this is a verifiable synthesis of *μ’* where each intermediate mutant is permissible. In other words,*μ’* is P-realizable, and this completes the proof of Part 2 as well as that of the theorem.⃞

In view of the above theorem, we can use the notion of C-realizability to develop a dynamic programming (DP) algorithm to solve the EVPS problem for a target mutant *μ’*. Let | *R’*| = *k* and let ⊆(R’) = 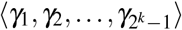 denote an ordered collection of all the non-empty subsets of where the subsets are in *non-decreasing* order of size; thus, for each i, 1 ≤ *i* ≤*k* — 1, all subsets of size i precede all those with size *i* + 1. The DP algorithm uses a Boolean array Β of size 2^k^ — 1. For convenience, we will index the elements of B with the subsets in *ℙ (R’*); that is, Β[*γ*] denotes the entry of the array corresponding to the subset *γ* ⊆ℙ (*R’*). (In the running time analysis presented in Section 4.6, we discuss a data structure to implement this scheme.) The significance of Β is as follows. For each *γ* ⊆ ℙ (*R’*), *Β*[*γ*] = TRUE if the mutant whose representative set is *γ* is permissible and C-realizable; otherwise, *Β*[*γ*] = FALSE.

Algorithm 3 shows how the entries of *Β* can be computed using DP. We can then determine whether *μ’* can be synthesized from the value of Β[*R*’]. The correctness of Algorithm 3 is a direct consequence of Theorem 4.4. In Section 4.6, we show that the algorithm can be implemented to run in *O*(*k* 2^2k^ +k *2^k^* (*p* + *s*)) time, which establishes that EVPS is FPT with respect to k. In Section 4.6, we also explain how to modify the algorithm to produce a permissible and verifiable synthesis with a minimum number of crosses; the running time and space requirements remain the same.

#### 4.6 Running Time Analysis and Extensions of Algorithm 3 for the EVPS Problem

Throughout this section, we will refer to the steps of the dynamic programming algorithm for the EVPS problem shown in Algorithm 3.

##### Analysis of running time and space used

We start with a discussion of a data structure that can be used in the implementation of the algorithm. Since | *R’*| = *k*, each subset of *R’* can be represented by a *k*-bit vector.

The Boolean array *Β* can be implemented as a *k*-level **binary trie** [14] where the leaves represent the array elements. The amount of memory used by this data structure is *O*(2*^k^*). Given the *k*-bit vector for a particular subset *γ* of *R’*, the leaf node corresponding to Β[*γ*] can be accessed in *O*(*k*) time by following the path from the root of the trie to the leaf node.

Using the trie data structure for the array *B*, Step 1 of the algorithm can be implemented to run in time *O*(*k*^2^). The number of iterations of the for loop of Step 2 is at most 2*^k^*. In each iteration, testing whether the subset *γ* is permissible (i.e whether it is in P) can be done in *O*(*kp*) time. Likewise, in each iteration, testing whether the subset *γ* represents a source mutant can be done in *O*(*ks*) time. Thus, over all the 2^k^ iterations of the loop in Step 2, these checks use *O*(2*^k^k*(*p* + *s*)) time.

Now, we estimate the time used in Step 2(d). In each iteration, Step 2(d) considers a subset *γ* of *R’*, with |*γ*| ≤ *k*. For such a subset *γ*, there are at most 2*^k-1^* partitions into two subsets. For any partition of *γ* into *γ*_1_ and we can determine the values of *Β* [*γ*_ι_] and *Β* [γ_2_] in *O*(*k*) time. Thus, the time for Step 2(d) in each iteration is *O*(2*^k^*-1 *k*). Since the number of iterations of the loop is at most 2^k^, the running time of Step 2(d) over all the iterations is *O*(2*^k^* 2*^k^*-1 *k*) = *O*(*k*2^2^*^k^*). Therefore, the total time for Step 2 is *O*(*k*2^2^*^k^* + *k* 2*^k^* (*p* + *s*)). Since Step 2 is the dominant part of the algorithm, the running time of the algorithm is also *O*(*k*2^2^*^k^* + *k*2*^k^(p* + *s*)). The algorithm uses *O*(2*^k^*) space for the array Β and *O*(*k*2*^k^*) space for the ordered set collection *P*(*R’*). Further, the representative set of each source and permissible mutant is at most *k*. So, the space for all the representative sets is *O(k(p* + *s*)). Since p and s are both at most 2*^k^*, the space requirement is dominated by that of the ordered collection ℙ (*R’*). Therefore, the space used by the algorithm is *O*(*k*2*^k^*).

##### Extension to obtain a synthesis with a minimum number of crosses

We can extend the dynamic programming algorithm so that whenever there is a permissible and verifiable synthesis, it finds one that uses the smallest number of crosses. To achieve this extension we use an enhanced version of array B. In this version, each element Β[*γ*] is a record with three fields which we denote by flag, min-crosses and partition. The flag field indicates whether *γ* can be synthesized in a permissible and verifiable manner. When Β[*γ*] .flag is TRUE, the min-crosses field stores the minimum number of cross operations needed to obtain a permissible and verifiable synthesis of *γ* and the partition field stores one partition of *γ* into two subsets *γ*_1_ and γ _2_ such that Β[*γ*_1_] = Β[*γ*_2_ = TRUE and the value Β[*γ*_1_].min-crosses + Β[*γ*_2_.min-crosses is the smallest among all the partitions of *γ*. Since the dynamic programming algorithm considers all partitions of any set *γ*, the values of the fields of each record Β[*γ*] can be suitably updated. At the end, if Β[*R*’] .flag is TRUE, then the value Β[*R*’] .min-crosses gives the minimum number of cross operations needed to obtain a synthesis of *μ’*. It is not difficult to see that the running time of and the space used by used by this version remain *O*(*k*2^2^*^k^* + *k*2*^k^*(*p* + *s*)) and O(*k*2*^k^*) respectively.

##### Extension to multiple mutants

There are two ways to extend the algorithm to synthesize multiple mutants. One way is to run the algorithm separately for each mutant. If t is the number of target mutants and k is the maximum size of the representative set of a target mutant, the running time of this approach is *O*(*t[k*2^2^*k* + *k*2*^k^*(*p* + *s*)]) and the space used is *O*(*k*2*^k^*). A second approach is useful when there is significant overlap among the representative sets of the target mutants. Let *R’* denote the union of all the representative sets of the t target mutants, and let | *R’*| = i. Then, we can run the algorithm once with the target mutant being the one whose representative set is *R_t_^’^*. From the solution array constructed by the algorithm, we can determine theanswerfor each of the target mutants. The running time of this approach is *O(ℓ* 2^2^*ℓ* + *ℓ* 2*^ℓ^* (*p* + *s*)) and the space used is O(i2^i^).

## 5 Experimental Results

As we noted in the introduction, there are no published databases that contain several mutants with three or more mutations that have been made experimentally. Therefore, we had to create novel datasets to test and evaluate our algorithms. We generate two complementary types of datasets: simulated and synthetic. Our evaluations on simulated datasets seek to replicate realistic scenarios faced by experimentalists who make new mutants from existing strains in order to validate computational predictions [7, 1, 15] (Section 5.1). To this end, we simulate mathematical models of the cell cycle in budding yeast in order to generate biologically interesting and meaningful target and permissible mutants. Synthetic datasets allow us to assess the variation in the performance of Algorithms 2 and 3 for the VS-FL and VPS problems as we vary the value of the corresponding fixed parameter (Section 5.2).

To generate the results for VS-FL, we used a 0-1 integer linear program (ILP) to solve the DCS instances (see Section 4.3). The ILP was easier to implement than the algorithm in [13]. We used the CPLEX software to solve the ILP. We noted that writing and reading files to interface between our software and CPLEX consumed a significant fraction of the running time. Therefore, we mention this cost separately when we report the running times of our algorithms. For the VPS problem, we implemented a recursive algorithm with caching, as an alternative to the DP. We used a computer with a 3.4GHz Intel i7 CPU and 16GB RAM running the 16.04 Ubuntu Linux operating system.

### 5.1 Evaluation on Cell Cycle Model Simulations

#### Simulations to Compute Target and Permissible Mutants

We selected a dynamic model of the budding yeast cell cycle [15] since we use its predictions in an ongoing collaboration on making and characterizing novel cell cycle mutants [15, 1, 23]. This model accurately simulates the phenotypic properties of more than 250 cell cycle mutants in budding yeast that have been published in the literature. Starting from 35 single-gene mutations, we simulated this model for mutations in all combinations of up-to-six genes. For each mutant, we recorded whether the simulation predicted the phenotype as “viable” or “inviable.”

Given two mutants *a* and *b* with respective representative sets *R_a_* and *R_b_*, we defined *b* to be a *parent* of *a* if | *R_a_ — R_b_*| = 1, i.e., a carries one more single gene mutation than *b*. We say that *b* is an *ancestor* of *a* if |*R_a_* — *R_b_*| | ≥ 1. We further define two specific classes of mutants, namely **rescued** and **synthetic lethal.** We define a mutant *a* to be *rescued* if *a* is predicted to be viable but had a parent b that is either experimentally known or predicted by the model to be inviable. We note that a rescued mutant may be *redundant*. For example,*^>Μ5Δ* rescues the inviable mutants *bck2Δcln3Δ* and *bck2Δcln3Δpds1Δ*. Since *pds1Δ* is itself a viable strain, the loss of Pds1 is irrelevant to the rescue phenotype. So, we consider only non-redundant rescued mutants, i.e., *bck2Δcln3Δwhi5Δ* but not *bck2Δcln3Δpds1Δwhi5Δ*. We define a mutant *a* to be *synthetic lethal* (abbreviated as SL) if *a* is inviable and all of a’s parents are viable.

Only 8% of the 1.6M mutants we simulated were viable (Table 1) The percentage of viable combinations decreased with an increase in the number of genes mutated, which supports our consideration of permissible mutants in the VPS problem. There were 77,243 rescued mutants of which 2,041 (2.6%) were non-redundant. Out of 1.48M inviable mutants, only 246 (0.016%) were SL. We specified inputs to our algorithms as follows:

**Table 1:** Statistics on simulations. We ignored combinations that contained redundant pairs of mutations, e.g., *cdh*1 Δ and CDH1 .constitutive. We abbreviate “number of mutations” as “#Ms,” “non-redundant” as “non-red,” and synthetic lethal mutants as “SL.”

| #Ms | Mutants | Viable | Rescued | Rescued non-red. | SL |
| --- | --- | --- | --- | --- | --- |
| 1 | 35 | 26 (74%) | - | - | - |
| 2 | 586 | 285 (49%) | 14 | 14 | 49 |
| 3 | 6,250 | 1,896 (30%) | 344 | 170 | 55 |
| 4 | 47,705 | 8,872 (18%) | 3,170 | 516 | 57 |
| 5 | 277,531 | 31,060 (11%) | 16,722 | 1,098 | 58 |
| 6 | 1,279,720 | 84,709 (7%) | 56,993 | 243 | 27 |
| **Total** | 1,611,827 | 126,848 (8%) | 77,243 | 2,041 | 246 |

#### Target mutants

Two sets: all four-, five-, and six-gene (i) non-redundant rescued mutants and (ii) SL mutants.

#### Source mutants

49 mutants available to our experimental collaborators [1] (Neil Adames and Jean Pec-coud, personal communication), including 35 single, 11 double, and three triple mutants (Table 3 in Section A.1 in the Appendix).

#### Permissible mutants

All viable mutants in our simulations.

#### Results for VS-FL

We used Algorithm 2 for VS-FL to compute the optimal synthesis for each target mutant. The total running time for these computations was 22.37 seconds (0.011 second per mutant), with an additional 91.92 seconds (0.045 second per mutant) incurred for file input and output related to the ILPs for the DCS problem. Figure 3(a) summarizes these results for rescued mutants and for SL mutants. The majority of target mutants required one fewer cross than the number of mutated genes (e.g., four crosses for a five-gene target mutant). The VS-FL algorithm capitalized on double mutants in the source set to compute syntheses with three crosses for 217 five-gene rescued mutants and syntheses with two crosses for 68 four-gene rescued mutants. For 26 five-gene mutants, the algorithm could compute syntheses with just two crosses, thus taking advantage of triple gene mutants in *S*. We observed similar trends for SLs.

**Figure 3:**
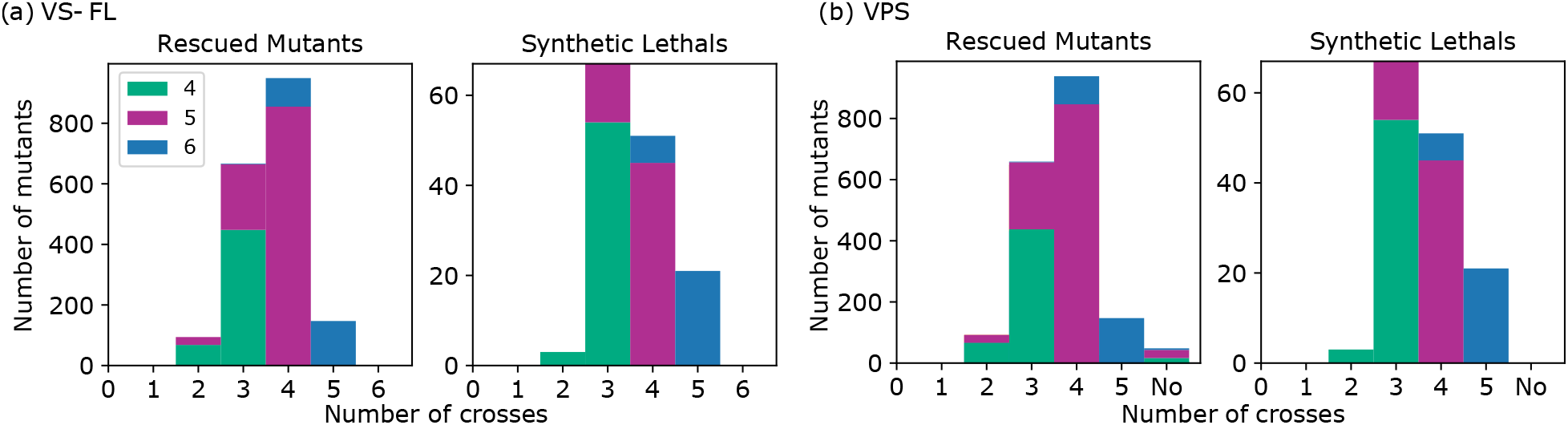
Distribution of the number of crosses in optimal syntheses for rescued mutants and for SL mutants (*x*-axis: number of crosses, y-axis: number of mutants with an optimal synthesis of that size, legend: the number of mutations in each target mutant). (a) Results for Algorithm 2 for VS-FL. (b) Results for Algorithm 3 for VPS. *x*-axis label of “No:” mutants without a verifiable, permissible syntheses.

#### Results for VPS

We used Algorithm 3 to compute an optimal synthesis for each target mutant in the presence of permissible mutants. The total running time for these computations was 18.5 seconds (average of 0.01 second per mutant). Figure 3(b) shows the results. We found that 1,992 of the 2,041 (97.6%) of the rescued mutants and all SL mutants had a verifiable, permissible synthesis. Of the 49 target mutants that did not have such a synthesis, there were 18 four-gene, 26 five-gene, and 5 six-gene mutants. Figure 4 shows such a mutant. In the caption of the figure, we explain why this mutant does not have a verifiable, permissible synthesis.

**Figure 4:**
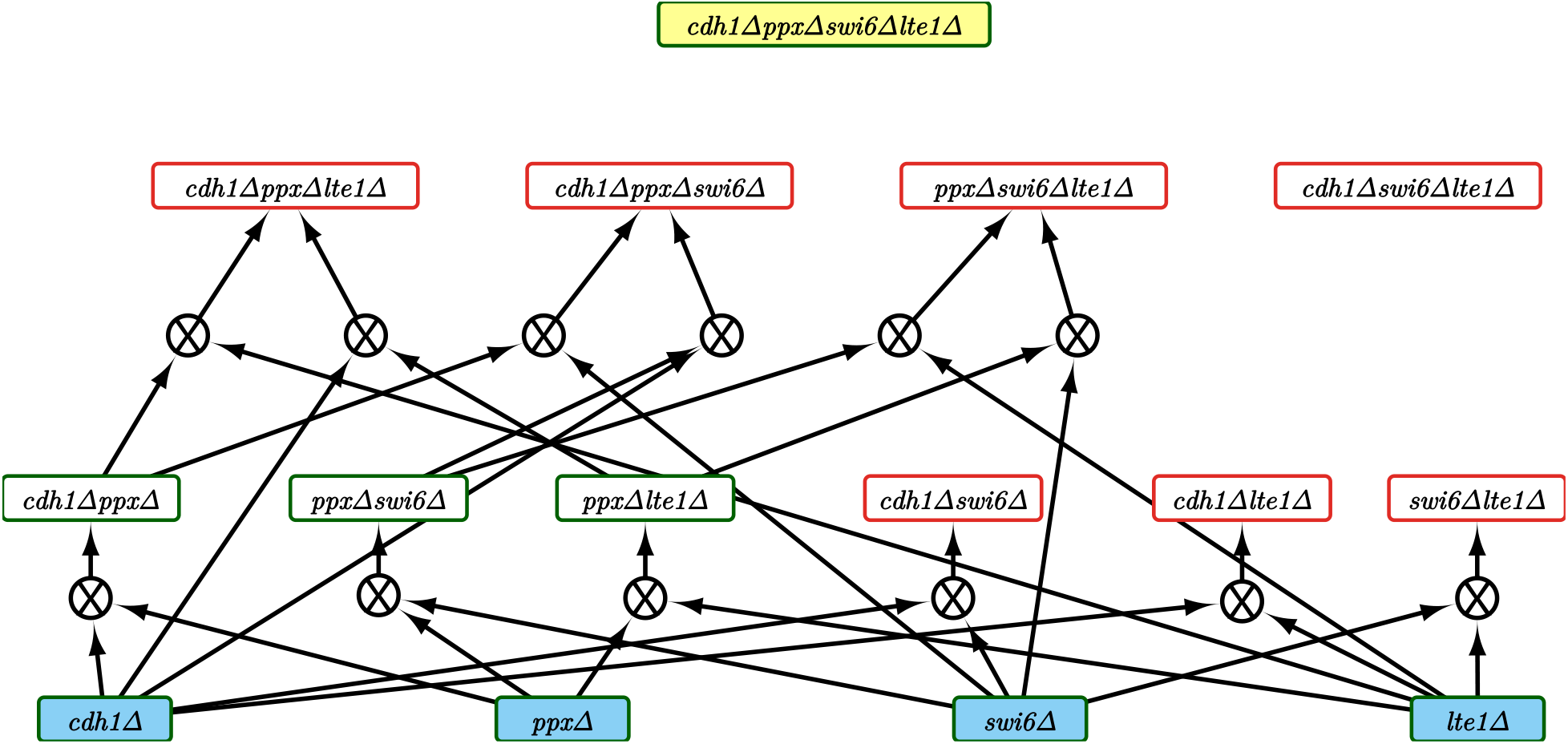
Example of a target rescued mutant for which there is no verifiable permissible synthesis. Each rectangular node is a mutant. A permissible mutant is in green (the simulations predict it to be viable). A non-permissible mutant is in red (the simulations predict it to be inviable). Further, source mutants are highlighted in blue, and the target mutant is highlighted in yellow.The label on a node lists the genes deleted in that mutant. Each blue circle represents a verifiable cross. Each such node has two incoming edges and one outgoing edge. This network contains all crosses involving the displayed mutants; it is identical to the genetic cross graph used by our ILP-based algorithm [25]. The node at the top of the network is the target rescued mutant. It does not have a permissible synthesis since (i) all its parents (triple mutants) are inviable and (ii) only one mutant is viable in every pair of double mutants that could be crossed in a verifiable synthesis of the target mutant. An alternative way of stating the second reason is that every pair of viable double mutants has one common gene, which means that crossing these double mutants is not a verifiable synthesis.

For each mutant with a verifiable, permissible synthesis, we compared the syntheses computed by the VS-FL and the VPS algorithms to determine if the restrictions imposed by permissible mutants increased the number of crosses in the synthesis. The size of the synthesis did not increase for most target mutants. However, four rescued mutants did have a larger synthesis for the VPS problem than for the VS-FL problem.

Figure 5(a) shows a synthesis with two crosses that is not permissible and Figure 5(b) shows a verifiable, permissible synthesis of the same mutant with three crosses.

**Figure 5:**
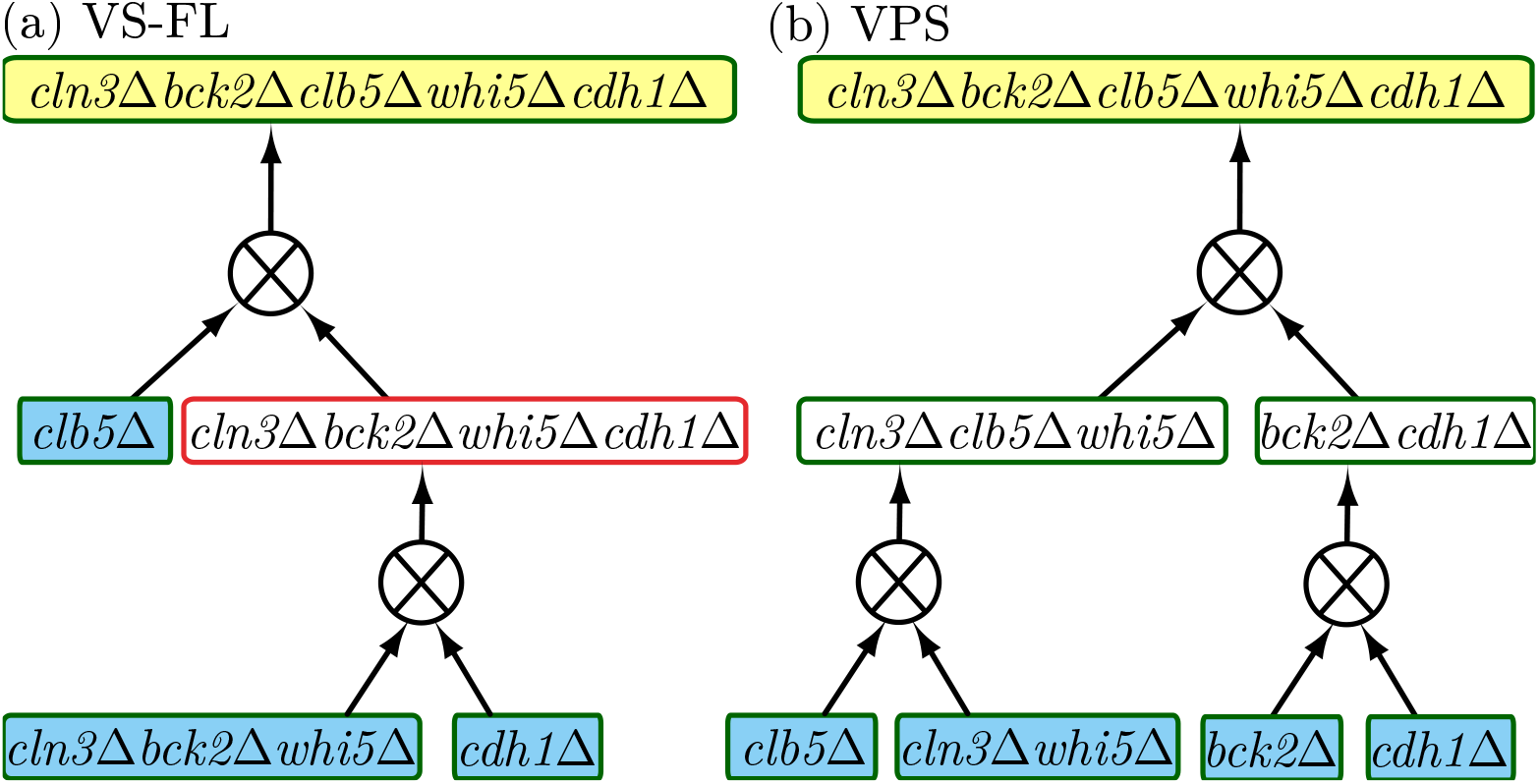
(a) Example of a verifiable but non-permissible synthesis with two crosses computed by Algorithm 2 for VS-FL. Node colors are as in Figure 4. (b) Example of a verifiable, permissible synthesis with three crosses computed by Algorithm 3 for VPS. Note that even though the triple mutant *cln3Abck2Awhi5A* is a source, Algorithm 3 does not use it in the synthesis since both *cln3Abck2Awhi5Acdh1A* and *cln3Abck2Awhi5Aclb5A* (not shown) are not permissible mutants (they are inviable).

These results show that permissible mutants can affect the synthesis size, even if only for a small number of target mutants. More importantly, the requirement of permissibility can preclude the existence of a synthesis for several target mutants. Our algorithms can handle the additional restrictions imposed by permissibility.

### 5.2 Evaluation on Synthetic Datasets

#### Generating Synthetic Data

Recall that Algorithm 2 for the VS-FL problem is FPT in *q*, the number of source mutants with three or more mutations, while Algorithm 3 for the VPS problem is FPT in *k,* the number of mutations in the target mutant. We created synthetic datasets that varied both parameters so that we could the same inputs to both algorithms. We created representative sets for all mutants from a universe *U* of 35 genes; we used this number to match the cell cycle model.

#### Target mutants

We generated a random target mutant whose representative set contained *k* = 6, 8, 10, or 12 genes from *U*. We used these values of *k* to test our algorithms on larger target mutants than we used in Section 5.1, where every mutant contained at most six mutations. For each of these four values, we report results averaged over 100 target mutants. Let *U*’ C *U* be the union of the representative sets of these 100 target mutants.

#### Source mutants

We started with *S* containing all singleton and doubleton subsets of *U*’. This choice ensured that each target mutant had a synthesis with at most *k/2* - 1 crosses, thereby making the results depend only on the triple mutants added. Next, we varied *q* from 0 to 300 in steps of 100 and added *q* triple mutants for S. In selecting the representative sets of these mutants, we sampled subsets of *U*’ uniformly at random. An experimentalist seeking to make a *k* -gene target mutant is likely to first make one or more triple mutants that are ancestors of the target mutant. We developed this strategy of selecting triple mutants to mimic the experimentalist. We stress that we use *U*’ only to select source mutants. We still apply our algorithms to each target mutant independently.

#### Permissible mutants

For each triple mutant *μ* we added to S, we considered every descendant of *μ* to be permissible, as long as the representative set of that descendant was a subset of *U*’.

#### Results for VS-FL

Figure 6(a) plots how the average number of crosses needed to make a target mutant changes with q. When *q* = 0, the algorithm for VS-FL runs Algorithm 1 for VS-2, and computes syntheses that only involve double mutants. As we increase *q,* the average number of crosses decreases as the synthesis can now take advantage of triple gene mutants. However, as the number of triple gene mutants in the source set increases, the time taken to run the algorithm for VS-FL increases considerably, especially for targets with 12 mutations (Figure 6(b)), as we may expect from the exponential dependence on *q* of the worst-case running time of Algorithm 2. For target mutants with 12 mutations, there is a considerable variation in the running times, especially for *q* = 300. Instead of plotting this error bar in Figure 6(b), we display the distribution of running times in Figure 6(c). While the algorithm ran in less than one second for over 75% of the target mutants, it took around two seconds for four mutants and over three seconds for two other mutants. Table 2 summarizes the time taken to read and write the ILPs for the DCS problem.

**Figure 6:**
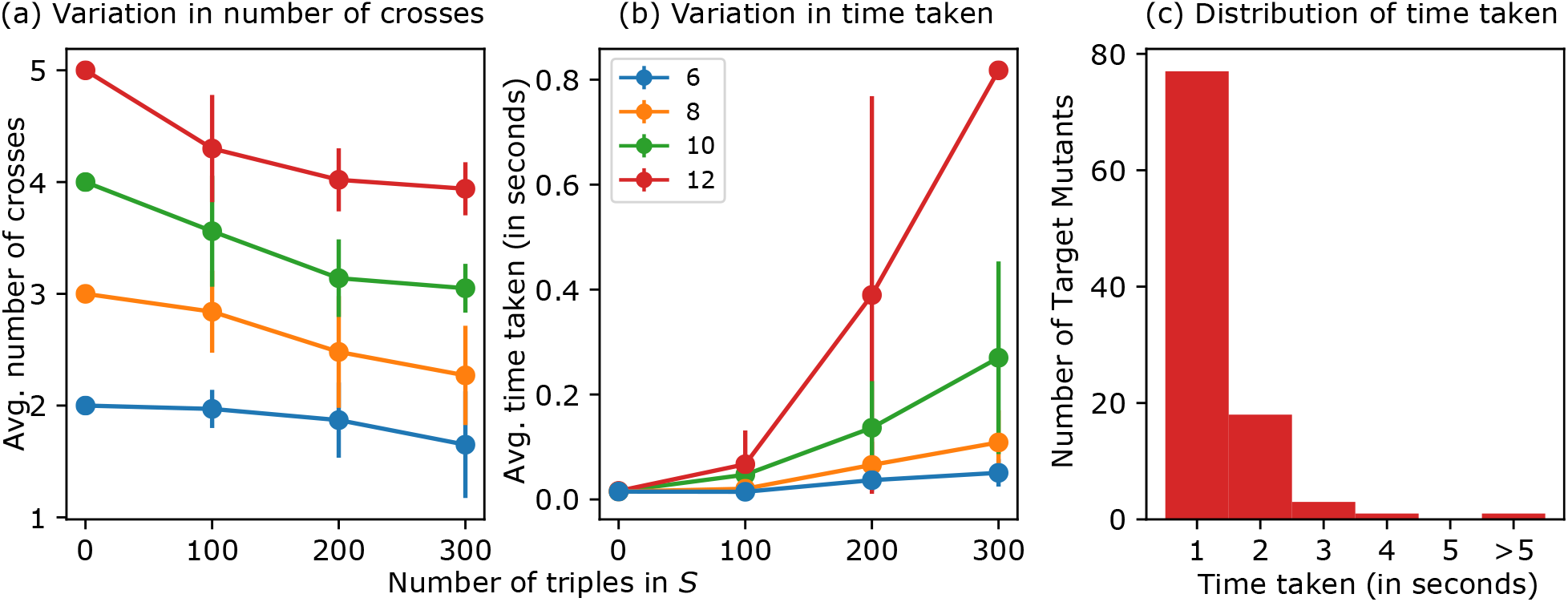
VS-FL results for synthetic data. Error bars indicate one standard deviation. The legend in (b) shows *k*, the size of the representative set of the target mutant. (a) The average number of crosses in an optimal synthesis as we change *q,* the number of triples in *S*. (b) Average time in seconds as we vary *q*. (c) Distribution of running time for 100 target mutants, *k* = 12, *q* = 300.

**Table 2:**
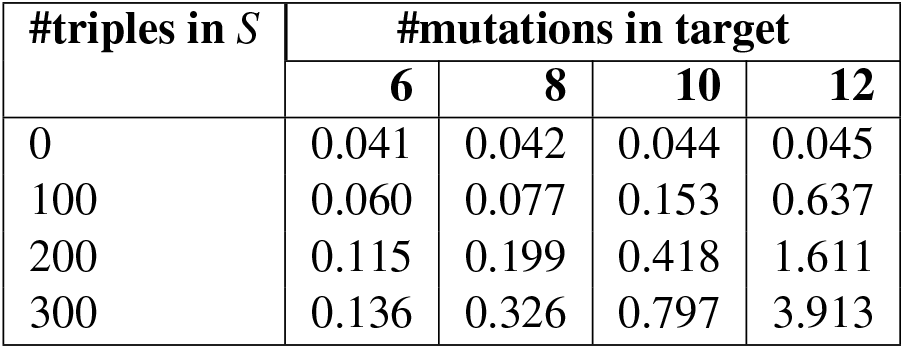
Average time taken for reading and writing the ILP for DCS problem for synthetic datasets.

| #triples in $S$ | #mutations in target | | | |
| --- | --- | --- | --- | --- |
|  | 6 | 8 | 10 | 12 |
| 0 | 0.041 | 0.042 | 0.044 | 0.045 |
| 100 | 0.060 | 0.077 | 0.153 | 0.637 |
| 200 | 0.115 | 0.199 | 0.418 | 1.611 |
| 300 | 0.136 | 0.326 | 0.797 | 3.913 |

#### Results for VPS

As we increase *q,* the number of triples in *S,* we also increase the size of the permissible set. Therefore, the number of crosses in an optimal synthesis decreases with an increase in *q* (Figure 7(a)). The average time taken by the VPS algorithm is negligible for *k* = 6,8,10 (Figure 7(b)), in contrast with the result for VS-FL (Figure 6(b)). For *k* = 12, the average time taken by the algorithm for VPS increases considerably compared to smaller values of *k* (Figure 7(b)), since the algorithm is FPT with respect to this parameter; we observe (but do no show here) a similar trend for *k* = 14. Moreover, the running time also increases with *q:* the more permissible mutants there are, the longer Algorithm 3 takes to find an optimal synthesis. Nevertheless, all running times are at most 20 seconds. Comparing Figure 7(b) and Figure 6(b) for *k* = 12 and *q* = 300, the VPS algorithm is about 18 times slower than the VS-FL algorithm.

**Figure 7:**
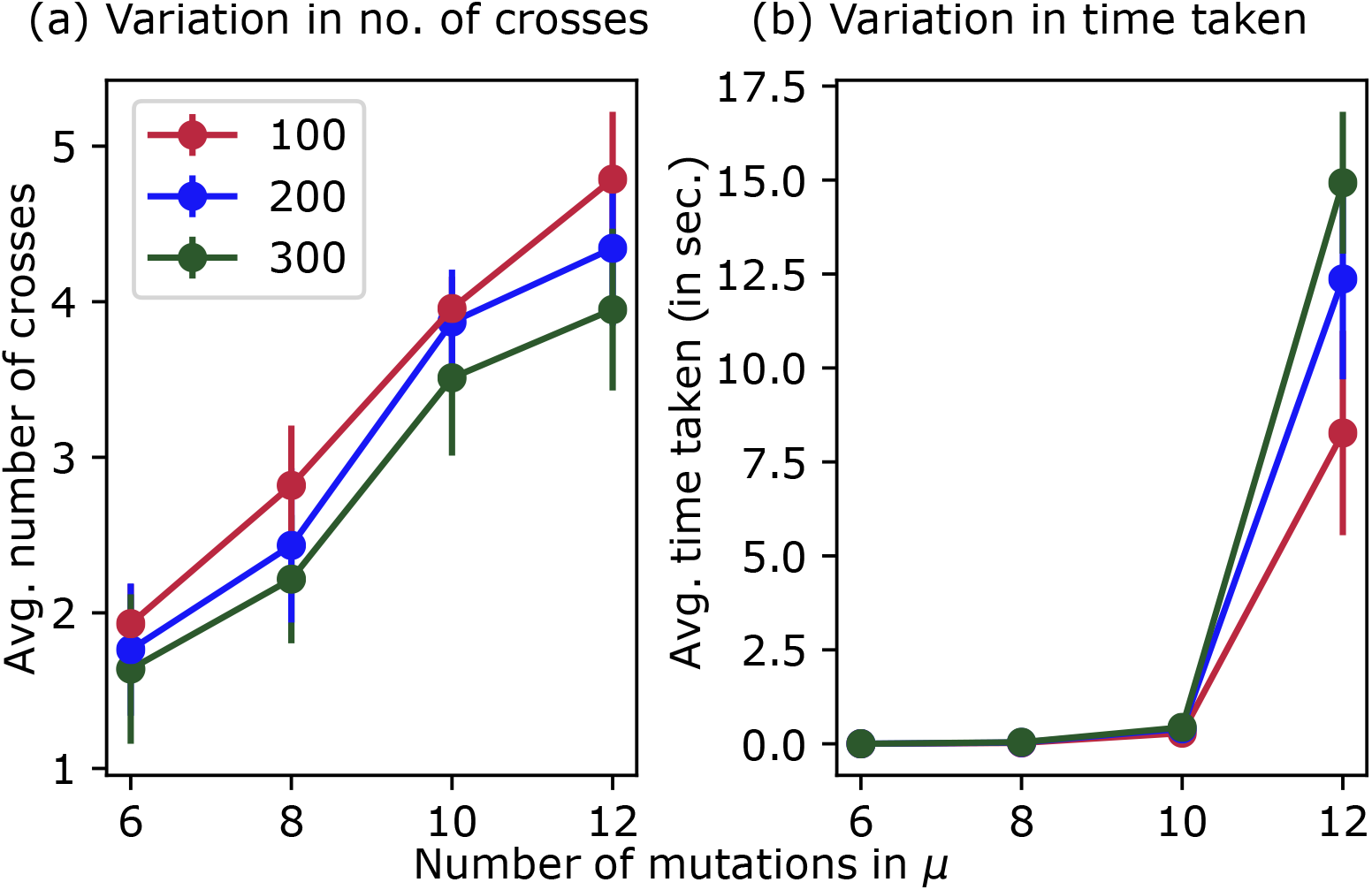
VPS results for synthetic data, averaged over 100 target mutants. Error bars indicate one standard deviation. The x-axis corresponds to *k*, the size of the representative set of the target mutant. The legend in (a) shows *q*, the number of triples in *S*. (a) Average number of crosses in an optimal synthesis as we vary *k*. (b) Average time taken in seconds.

#### Comparing VPS to CrossPlan

Here, we compare the VPS algorithm to our ILP-based approach, which we call CrossPlan [25]. We run both algorithms to compute an optimal verifiable synthesis for each target mutant with *k* = 12. For a specific target mutant *μ*, in addition to the inputs to VPS, CrossPlan takes two additional inputs: (a) l, an upper bound on the number of crosses needed and (b) a genetic cross graph G^). We set *l = k/2 —* 1, since *S* contains all double mutants. The graph (G^) contains one *mutant node* for each permissible mutant whose representative set is a subset of *μ’* s set. For every pair of permissible mutants with disjoint representative sets, (G^) contains a *cross node* that represents the corresponding genetic cross. In addition, (G^) contains two incoming edges and one outgoing edge for each cross. See Figure 4 for an example. We compute this graph independently for each target mutant. On average, a graph contains 4,095 = 2^12^ − 1 mutant nodes and 66,000 cross nodes.

Both algorithms report optimal synthesis of the same size or that no verifiable, permissible synthesis exists. However, the VPS algorithm is over 28 times faster (24.83 minutes vs 700.77 minutes) in computing optimal syntheses. It was 10 times faster than CrossPlan (9 minutes vs. 90.08 minutes) for the question of determining if a verifiable, permissible synthesis exists. The CrossPlan ILP contained 300K variables and 600K constraints, on average. In fact, due to the size of the ILP, we set a time limit of six minutes on the Gurobi solver for each target mutant since in our experience the solver computes a heuristic solution quickly but can spend hours in proving its optimality. These results show the benefit of the strategy we have adopted in this paper of formulating synthesis problems that do not need the explicit construction of the genetic cross graph.

## 6 Conclusions

We introduced a fundamental problem in computational biology: how to use genetic crosses to efficiently synthesize a target mutant from source mutants. We formalized this question in several ways that incorporated key experimental considerations of verifiability and permissibility. We showed that checking the existence of a synthesis is **NP** -complete. We presented one polynomial time and two FPT algorithms for these problems. On simulated and synthetic data, these algorithms ran efficiently and provided useful results.

There are several interesting directions for future research. Developing FPT algorithms that compute an optimal synthesis for multiple target mutants is an important open problem. These methods will complement the ILP-based solution we have proposed [25].

It may also be useful to study the scenario when permissible intermediate mutants are specified implicitly rather than explicitly. A simple example of an implicit specification is to allow every intermediate mutant to be permissible, which is identical to the VS problem. Another example is the following: an intermediate mutant is permissible if and only if it has no gene from the set {g_1_, g_2_, g_3_}; this set could correspond to genes whose products form an essential protein complex. Such a description can specify an exponentially large set of permissible intermediate mutants. However, given a target mutant *μ*, we can efficiently determine whether or not it is permissible under the given specification. Our algorithm for VPS in Section 4.5 can be readily modified to work for such implicit specifications of permissible intermediate mutants.

Our current model for creating new mutants includes only the genetic cross. It is important to extend this model to include other experimental techniques, especially genome editing methods, while considering the relative costs of each approach. We are also interested in studying and incorporating more complex constraints imposed by genetic crosses in model organisms.

## Footnotes

1 This notation represent the deletion of the Whi5 gene.

## Acknowledgments

National Science Foundation grants CCF-1617678, ACI-1443054, and IIS-1633028 supported this work. We also thank the reviewers for their feedback.

## A Appendix

### A.1 List of Source Mutants

Table 3 shows the 49 source mutants we use in the evaluation using simulations of the cell cycle model (Section 5.1).

**Table 3:** The 49 source mutants used in Sec. 5.1. CDH1 .constitutive is a mutant that expresses CDH1 constitutively. CLB1 *clblA* refers to a strain with *clb2* deletion with wildtype CLB1 (CLB1 and CLB2 are paralogs that are redundant in function). *apc^ts^* refers to a non-functional Anaphase-promoting Complex, which is achieved by using a temperature sensitive strain. TAB6-1 refers to *telophase arrest bypass* mutants representing altered Cdc14:Net1 stoichiometries at mitotic exit.

| Single gene mutants | Double gene mutants |
| --- | --- |
| *cln3*Δ | *cln3*Δ*bck2*Δ |
| *bck2*Δ | *cln3*Δ*whi5*Δ |
| *sic1*Δ | *cln3*Δ*mbp1*Δ |
| *cki*Δ | *bck2*Δ*whi5*Δ |
| *cln2*Δ | *bck2*Δ*mbp1*Δ |
| *whi5*Δ | *cln2*Δ*whi5*Δ |
| *cdh1*Δ | *cln2*Δ*mbp1*Δ |
| CDH1_constitutive | *whi5*Δ*swi4*Δ |
| *clb2*Δ | *whi5*Δ*mbp1*Δ |
| CLB1 *clb2*Δ | *whi5*Δ*swi6*Δ |
| *cdc20*Δ | *mbp1*Δ*swi6*Δ |
| *clb5*Δ |  |
| *swi5*Δ | **Triple gene mutants** |
| *apc<sup>ts</sup>* | *cln3*Δ*bck2*Δ*whi5*Δ |
| APCA | *cln3*Δ*whi5*Δ*mbp1*Δ |
| *tem1*Δ | *cln3*Δ*mbp1*Δ*swi6*Δ |
| *cdc15*Δ |  |
| *net1<sup>ts</sup>* |  |
| *cdc14<sup>ts</sup>* |  |
| TAB6–1 |  |
| *pds1*Δ |  |
| *ppx*Δ |  |
| *mad2*Δ |  |
| *bub2*Δ |  |
| *swi4*Δ |  |
| *mbp1*Δ |  |
| *clb5*Δ |  |
| *swi6*Δ |  |
| *cdc5*Δ |  |
| *cdc6*Δ |  |
| *cdc14–1*Δ |  |
| *ssa1*Δ |  |
| *ydj1*Δ |  |
| *lte1*Δ |  |
| *swe1*Δ |  |

